# The FACT complex is required for DNA demethylation at heterochromatin during reproduction in Arabidopsis

**DOI:** 10.1101/187674

**Authors:** Jennifer M. Frost, M. Yvonne Kim, Guen-Tae Park, Ping-Hung Hsieh, Miyuki Nakamura, Samuel Lin, Hyunjin Yoo, Jaemyung Choi, Yoko Ikeda, Tetsu Kinoshita, Yeonhee Choi, Daniel Zilberman, Robert L. Fischer

**Author notes:** To whom correspondence should be addressed E-mail addresses.

## Abstract

The DEMETER (DME) DNA glycosylase catalyzes genome-wide DNA demethylation and is required for endosperm genomic imprinting and embryo viability. Targets of DME-mediated DNA demethylation reside in small, euchromatic, AT-rich transposons and at the boundaries of large transposons, but how DME interacts with these diverse chromatin states is unknown. The STRUCTURE SPECIFIC RECOGNITION PROTEIN 1 (SSRP1), subunit of the chromatin remodeler FAcilitates Chromatin Transactions (FACT), was previously shown to be involved in the DME-dependent regulation of genomic imprinting in Arabidopsis endosperm. Therefore, to investigate the interaction between DME and chromatin, we focused on the activity of the two FACT subunits, SSRP1 and SUPPRESSOR of TY16 (SPT16), during reproduction in Arabidopsis. We find that FACT co-localizes with nuclear DME in vivo, and that DME has two classes of target sites, the first being euchromatic and accessible to DME, but the second, representing over half of DME targets, requiring the action of FACT for DME-mediated DNA demethylation genome-wide. Our results show that the FACT-dependent DME targets are GC-rich heterochromatin domains with high nucleosome occupancy enriched with H3K9me2 and H3K27me1. Further, we demonstrate that heterochromatin-associated linker histone H1 specifically mediates the requirement for FACT at a subset of DME-target loci. Overall, our results demonstrate that FACT is required for DME targeting by facilitating its access to heterochromatin.

## Introduction

Cytosine methylation regulates gene expression and silences transposable elements (TEs) in plants and vertebrates (1). In *Arabidopsis thaliana*, distinct DNA methyltransferases and pathways are responsible for establishing and maintaining DNA methylation in three sequence contexts: CG, CHG, and CHH, where H corresponds to A, T, or C (2). Gene body methylation is primarily CG, whereas transposable elements display methylation in all sequence contexts (3–5). Removal of DNA methylation occurs via the Base Excision Repair (BER) pathway, where dual function glycosylase/AP lyases catalyze excision of 5-methylcytosine from DNA and nick the sugar-phosphate backbone.

Downstream, AP-endonuclease, DNA polymerase and DNA ligase function to insert cytosine in place of the excised 5-methylcytosine (6). Demethylation of TEs that overlap gene regulatory regions influences gene expression: demethylation of transcriptional start sites and sequences that allow the binding of activating factors can promote expression, whereas demethylation of sequences that allow the binding of repressive factors can suppress gene activity.

Epigenetic reprogramming by DNA demethylation is vital for reproduction in mammals and flowering plants (7, 8). Flowering plants are the most evolutionarily successful and diverse group of plants on earth, and the defining feature of their reproduction is double fertilization. Double fertilization is mediated by multicellular male and female gametophytes, generated from haploid spores by multiple rounds of mitosis. The male gametophyte consists of two sperm cell nuclei and a vegetative cell nucleus, encased within the vegetative cell. A pollen tube germinates from the vegetative cell, delivering two sperm cells to the female gametophyte, where one fertilizes the haploid egg, which develops into the embryo, and the other fertilizes the homodiploid central cell, to form the triploid placenta-like endosperm. The embryo and endosperm, surrounded by maternal cell layers, comprise the seed. The vegetative and central cells, adjacent to the sperm and egg cells, respectively, are so-called gamete companion cells. In *Arabidopsis thaliana*, active DNA demethylation by the DNA glycosylase DEMETER (DME) occurs specifically in gamete companion cells, whereby highly specific transcriptional regulation during gametogenesis ensures DME expression is confined to these cells (9–11). DME-mediated DNA demethylation occurs at thousands of discrete loci genome-wide, including regulatory regions for genes encoding components of the Polycomb Repressive Complex 2 (PRC2); FERTILIZATION INDEPENDENT SEED 2 (FIS2) and MEDEA (MEA) inducing their monoallelic expression, i.e. genomic imprinting, in the endosperm (12). PRC2 confers H3K27me3 modifications that regulate gene expression and genomic imprinting during seed development. Activation of PRC2 component expression by DME is required for endosperm cellularization, a process essential to viable seed formation (13–15). Thus, maternal demethylation, initiated in the central cell, (9, 10) is vital for Arabidopsis reproduction, and loss of maternal DME results in seed abortion (9, 16–18).

Methylation removal by DME is catalyzed efficiently at CG, CHG and CHH (9, 17). DME acts in a targeted manner, and tends to demethylate relatively euchromatic TEs that are small, AT-rich, nucleosome-poor, and generally interspersed with genes in chromosome arms (9). DME also acts on longer heterochromatic TEs, primarily at their edges, and these TEs are prevalent in pericentromeric, gene poor regions, enriched with heterochromatic histone marks (9). How DME can successfully access regions of differing chromatin structure is not known. Chromatin structure is dictated by the organization of its functional unit, the nucleosome, consisting of an octameric core and often a linker molecule, histone H1. The core consists of two copies each of histone subunit pairs H2A/H2B and H3/H4, which can be further modified by posttranslational modifications of their NH_2_ terminal amino acids (19). Chromatin structure can also be altered through changes in nucleosome presence and spacing, and the exchange of canonical histone subunits for variant proteins, as catalyzed by ATP-dependent chromatin remodeling complexes and histone chaperones (19). FAcilitates Chromatin Transactions (FACT) is an essential, multi-domain protein complex conserved in eukaryotes, and capable of multiple interactions with nucleosome components, binding free H2A/H2B and H3/H4 dimers, as well as intact nucleosomes (20, 21). FACT is required for transcription initiation and elongation, nucleosome disassembly and reassembly - including histone variant exchange, notably of H2A.X; and for chaperoning free histones (20–22). In vertebrates and plants, FACT is a heterodimer of High Mobility Group (HMG) domain STRUCTURE SPECIFIC RECOGNITION PROTEIN 1 (SSRP1) and SUPPRESSOR of TY16 (SPT16) (23–25). Further supporting the role of chromatin in DME function, the smaller FACT subunit SSRP1 was previously shown to be involved in DME-mediated DNA demethylation at selected imprinted genes in Arabidopsis (26).

Here, we used Arabidopsis FACT complex mutants and analyzed DNA methylation genome-wide in developing seeds and the male gametophyte to delineate how chromatin structure affects DME targeting in Arabidopsis. We find that DME requires histone chaperone FACT to demethylate over half of its targets in the central cell. DME and the FACT complex protein SPT16 are located closely in the nucleus, and they interact, either directly or through local intermediates. We show that chromatin structure plays an important role in determining the degree to which FACT is needed for DNA demethylation. In regions with an elevated GC ratio, high nucleosome occupancy, and enriched for heterochromatin markers such as H3K27me1 and H3K9me2, FACT is required for DME access and activity. Moreover, we demonstrate that linker histone H1 mediates the requirement for FACT at a subset of DME-target loci.

## Results

### FACT is required for DME-mediated genome-wide DNA demethylation in the maternal endosperm

To assess the contribution of FACT to DNA demethylation during Arabidopsis reproduction, we analyzed DNA methylation in plants with a nonsense mutation in the gene encoding the small subunit of FACT, SSRP1; *ssrp1-3* (Figure 1A; (26)). FACT is required ubiquitously in eukaryotic cells for fundamental processes, including transcriptional elongation, and seeds homozygous for complete loss-of-function mutant alleles are not viable (26–28), therefore *ssrp1-3* mutant plants are always heterozygous.

**Figure 1.**
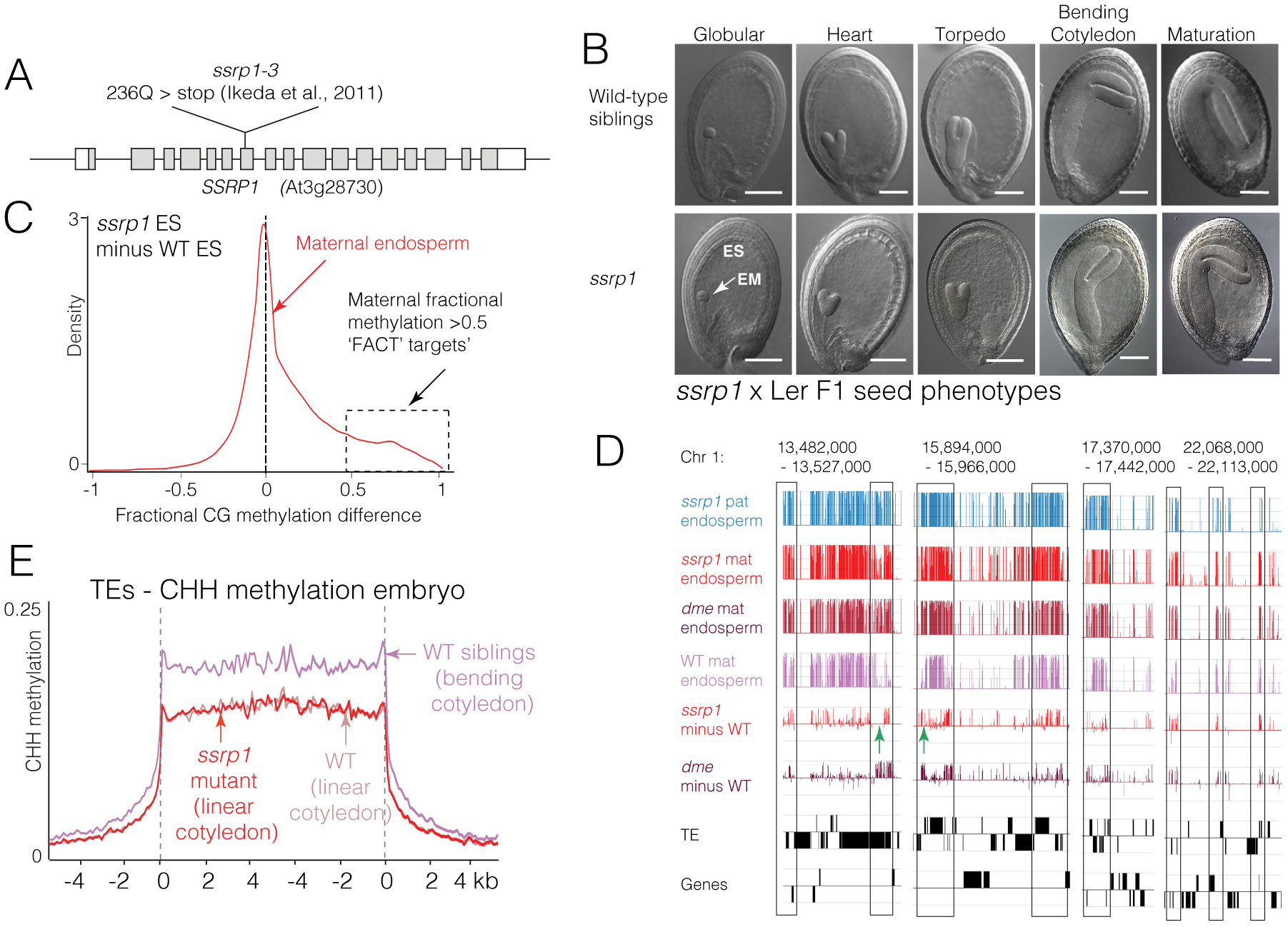
FACT is required for DME-mediated genome-wide DNA demethylation in the maternal endosperm. (A) Diagram showing the SSRP1 gene structure and location of the nonsense mutation *ssrp1-3* (26). (B) Photographs of developing F1 seeds from heterozygous *ssrp1-3* mutant plants crossed as females to wild-type L*er* pollen. ‘Delayed’ and ‘Normal’ seed fractions had respectively inherited mutant *ssrp1-3* and wild-type (WT) maternal alleles. The morphological stages of wild-type embryos are indicated. Scale bar = 100 um (C) Kernel density plot of CG methylation differences between *ssrp1-3* mutant endosperm (ES) and WT endosperm for the maternal allele. Positive numbers indicate hypermethylation and loci whose fractional methylation level is 0.5 or greater, i.e. SSRP1/FACT targets, are indicated by the dotted black box. (D) Genome browser alignments of DNA methylation with genes and TE annotations at selected loci of Arabidopsis Chromosome 1. Traces show raw CG DNA methylation scores for *ssrp1-3* paternal and maternal endosperm, *dme-2* maternal endosperm (9), and CG methylation differences for mutant minus WT in *ssrp1-3* and *dme-2* maternal endosperm genomes. Positive numbers indicate hypermethylation in the mutant genome. Regions of hypermethylation that overlap in each mutant are boxed. (E) Average CHH methylation in WT and *ssrp1-3* mutant developing embryos, aligned according to the 5’ and 3’ ends of TEs. WT sibling and ssrp1-3 embryos were from the same siliques at 9 DAP, but WT (linear cotyledon) embryos were dissected from siliques crossed at the same time, but embryos were taken at 7 DAP, to match *ssrp1-3* embryo development (i.e. linear cotyledon) at 9 DAP.

To analyze DNA methylation in the *ssrp1-3* mutant female gametophyte, we used developing endosperm as a proxy for central cells, where DNA demethylation takes place (10). Plants heterozygous for *ssrp1-3* in the Columbia-0 (Col-0) ecotype were pollinated with wild-type pollen from Landsberg erecta (L*er*) ecotype plants, and hand-microdissected F1 embryo and endosperm were isolated 8-10 days after pollination (DAP). The Col-0 and L*er* ecotypes differ by over 400,000 single-nucleotide polymorphisms (SNPs) that allow us to distinguish maternal and paternal genomes in F1 progeny (9). Whereas seeds from wild-type plants develop at the same rate within a given silique, and rarely abort, F1 siliques from heterozygous *ssrp1-3* plants crossed to L*er* contain three groups of seeds; approximately equal numbers of normally developing viable seed and delayed viable seed, (Figure 1B) and rare aborting seeds with uncellularized endosperm (26). We established the frequency of the *ssrp1-3* mutant (T) and WT (C) alleles in viable seeds by subcloning and DNA sequencing genotyping amplicons. Delayed seeds had maternally inherited the *ssrp1-3* mutant allele, with a 2:1 maternal to paternal ratio expected for the triploid endosperm, (130:63 T:C, c^2^ = 0.0117, p = 0.914) and normal seeds had only inherited the wild-type allele, (92 alleles counted, all C).

Next-generation bisulfite sequencing of endosperm from genotyped delayed (maternal mutant) and normally-developing (WT control) seeds revealed that the *ssrp1-3* maternal endosperm genome was hypermethylated genome-wide compared to wild-type in the CG context (positive shoulder in Figure 1C). As a control, we used developing embryos from normal and delayed seeds as a proxy to analyze DNA methylation in the egg. Methylation of both maternal and paternal alleles in delayed embryos (Figure S1A) and the paternal endosperm genome (Figure S1B) were identical to wild-type, indicating that hypermethylation was inherited specifically from the maternal mutant central cell.

DME activity in the central cell promotes endosperm demethylation of maternal DNA genome-wide, and was shown previously to require SSRP1 at certain sites (9, 10, 18, 26). Sites of maternal hypermethylation in the *ssrp1-3* mutant endosperm genome overlap with those in *dme-2* mutant endosperm (Figure 1D, black boxes); (9)), although regions hypermethylated only in *dme-2* mutant maternal endosperm are visible (Figure 1D, green arrows). CG hypermethylated loci in *ssrp1-3* endosperm were also maternally hypermethylated at CHG and CHH contexts, (Figure S1C and S1D), indicating that direct DME-mediated demethylation of CG, CHG and CHH cytosine contexts in the Arabidopsis central cell is dependent on FACT. In addition, genome-wide non-CG methylation was slightly reduced globally (Figure S1C and D) and at TEs (Figure S1E and F), likely due indirectly to DME promotion of PRC2 activity which, in turn, promotes non-CG DNA methylation (9, 29).

In the embryo, CG and CHG methylation were identical to WT (Figure S1A and S1G). However, we found that CHH methylation in TEs in embryo was lower in *ssrp1-3* mutants compared to WT (Figure 1E). Previously, Arabidopsis embryos were shown to undergo a developmental increase in CHH methylation (30), and at the time of measurement *ssrp1-3* mutant embryos were at the linear cotyledon stage compared to the bending cotyledon stage of WT sibling embryos (Figure 1B). We therefore re-measured CHH methylation in WT linear cotyledon embryos and found their CHH methylation levels to be identical to the *ssrp1-3* mutant linear cotyledon embryos (Figure 1E). This demonstrated that differential CHH methylation in embryos was due to developmental stage rather than the *ssrp1-3* mutation. Thus, we detected no direct effect on DNA demethylation in the Arabidopsis egg cell that is dependent on FACT.

### FACT is required for demethylation at >50 % DME DMRs in endosperm

To establish the extent to which FACT is required for DME-mediated DNA demethylation, we plotted the methylation status of *ssrp1-3* hypermethylated loci (fractional methylation difference compared to wild-type of >0.5 shown in Figure 1C) in *dme-2* mutant endosperm (Figure 2A, purple trace; (9)). This comparison resulted in a positive density peak, also >0.5 fractional methylation, showing that loci that become hypermethylated in *ssrp1-3* endosperm are also hypermethylated in *dme-2* mutant endosperm, and to a similar extent. A reciprocal analysis of *dme-2* hypermethylated loci in *ssrp1-3* mutant endosperm (Figure 2A, red trace) has two peaks; the positive peak at >0.5 fractional methylation represents loci that are hypermethylated in both *ssrp1-3* and *dme-2* mutants, i.e. they are shared DME and FACT target sites. Conversely, the peak centered on zero is indicative of sites that are only hypermethylated in *dme-2* mutant endosperm, i.e. they are targets of DME, but not FACT. By merging 50 bp windows within 300 bp regions that demonstrate a statistically significant difference in CG DNA methylation (Fisher exact test p<10^-3^), we created a set of FACT-mediated differentially methylated loci. If differential methylation over the whole region was significant (p<10^-10^), we defined the region as a FACT DMR, and identified 5186 FACT DMRs of at least 100bp, compared to 9816 DME DMRs defined previously (Figure 2B and Table S1; (9)). All 5168 FACT DMRs are a subset of DME DMRs. Thus, approximately 53 % of DME target sites in the endosperm require FACT for DNA demethylation, referred to hereafter as FACT-DME loci. 4648 DMRs do not require FACT for efficient DNA demethylation, referred to hereafter as DME-only loci. Thus, FACT is required for demethylation at over half the DME sites in the central cell.

**Figure 2.**
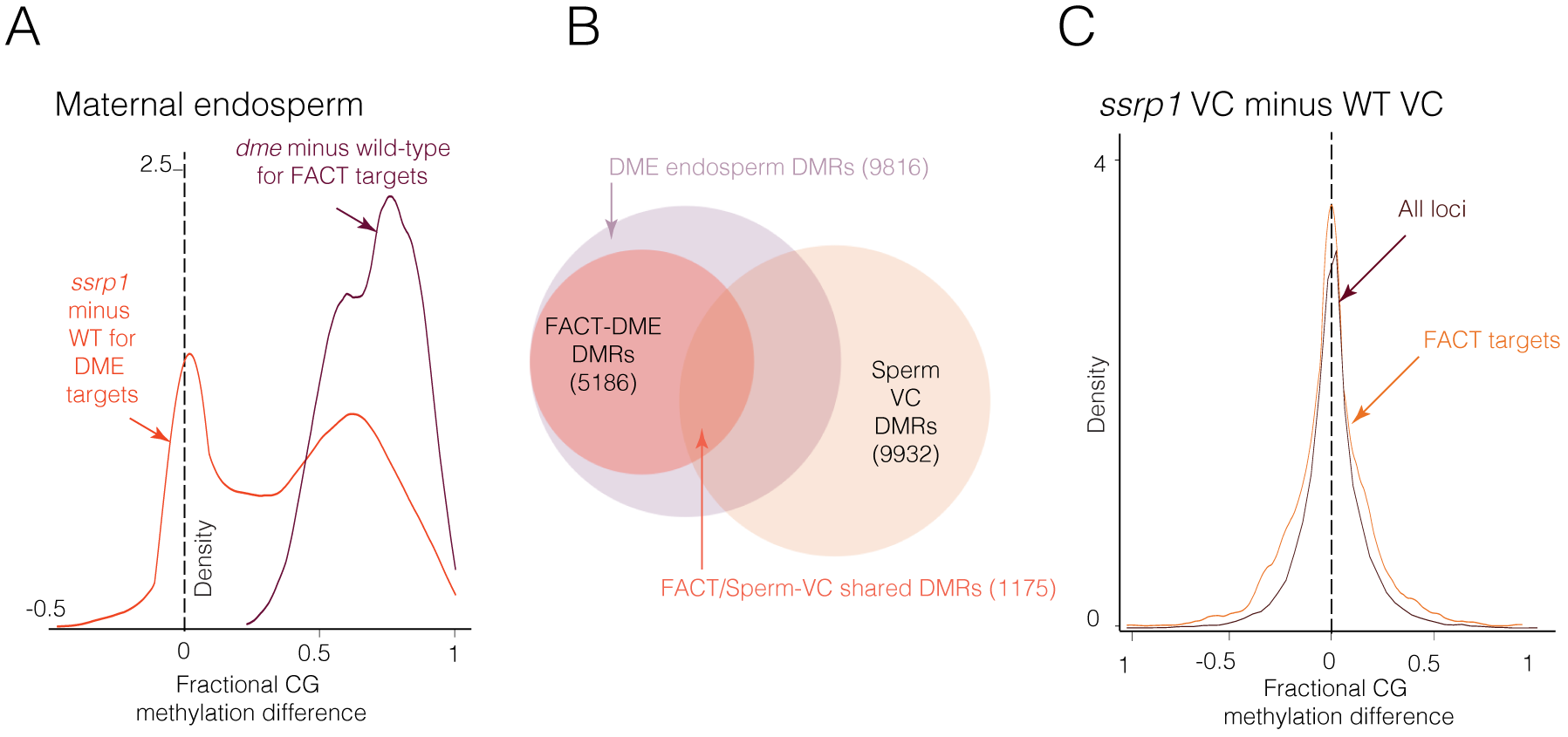
FACT is required for demethylation at >50 % DME DMRs in endosperm. (A) Red trace: Kernel density plot of CG methylation differences between *ssrp1-3* and wild type maternal genomes, specifically for DME targets, as defined by loci with fractional methylation > 0.5 in *dme-2* mutant endosperm compared to wild-type. Purple trace: Kernel density plot of CG methylation differences between *dme-2* and wild type maternal genomes, specifically for FACT targets, as defined above. (B) Venn diagram depicting the overlap between significant (Fisher’s exact test, P <10^-10^) differentially methylated regions (DMRs) of at least 100 bp in endosperm resulting from DME activity on the maternal genome (in the central cell), FACT activity on the maternal genome (FACT DMRs) and DME activity in vegetative cells (VC; Sperm/VC DMRs)). DME-only DMRs are those loci in endosperm which are targeted by DME but not FACT (C) Kernel density plots of CG methylation differences between *ssrp1* vegetative cells (VC) and WT VC, showing all loci (brown trace), and FACT CG target loci (orange trace) in maternal endosperm as defined above.

### We do not detect a requirement for FACT in pollen DNA demethylation

There are 9932 DME DMRs present between the sperm and vegetative cell genomes, and 1175 overlap with the 5186 DME DMRs in endosperm that are shared with FACT (Figure 2B and Table S1; (9)). We investigated whether FACT is also required for genome demethylation in the male gametophyte by isolating sperm and vegetative cell nuclei from fluorescence-activated cell sorted (FACS) pollen harvested from *ssrp1-3* heterozygous plants. Previous work shows that the *ssrp1-3* allele has a low rate of paternal transmission to F1 seeds (26). To establish whether the *ssrp1-3* mutant allele was present at a normal level in pollen, we cloned *ssrp1-3* genotyping amplicons in our pollen sample isolated from heterozygous *ssrp1-3* plants, as we did for endosperm, finding that the paternal *ssrp1-3* and wild-type alleles are present in pollen at approximately equal frequency (37:44, WT:*ssrp1-3* mutant, 0.84:1, c^2^ = 0.3012, p = 0.583). Thus *ssrp1-3* is transmitted normally through male meiosis and subsequent mitosis so that *ssrp1-3* mutant pollen is formed. We therefore suggest that the *ssrp1-3* male transmission defect identified by Ikeda and colleagues manifests after pollen formation (26). Bisulfite sequencing did not reveal hypermethylation of the mutant vegetative cell genome, either genome-wide or by specifically focusing on FACT target loci in endosperm (Figure 2C), and we did not identify any statistically significant DMRs between wild-type and *ssrp1-3* mutant vegetative cells. Thus, inheriting a mutant *ssrp1-3* allele does not seem to affect patterns of DME-mediated DNA demethylation in the vegetative cell.

### SPT16 contributes to DNA demethylation, and colocalizes with DME in nuclei

FACT consists of SSRP1 and the larger SPT16 subunit. To establish whether SPT16 mutants exhibited similar phenotypes to *ssrp1-3*, we obtained seeds carrying a T-DNA insertion in the coding region of *SPT16*, henceforth referred to as *spt16-3* (Figure 3A; Table S2 (31)). The seed phenotypes of heterozygous F1 *spt16-3* selfed or crossed to L*er* were very similar to *ssrp1-3,* with approximately the same ratios of normal and delayed seed development present (Figure 3B, S3A, Table S2 and S3), and plants could not be made homozygous. Delayed seeds were highly enriched for the *spt16-3* mutant allele (Figure S3B), although seed abortion in *spt16-3* was not above background.

**Figure 3.**
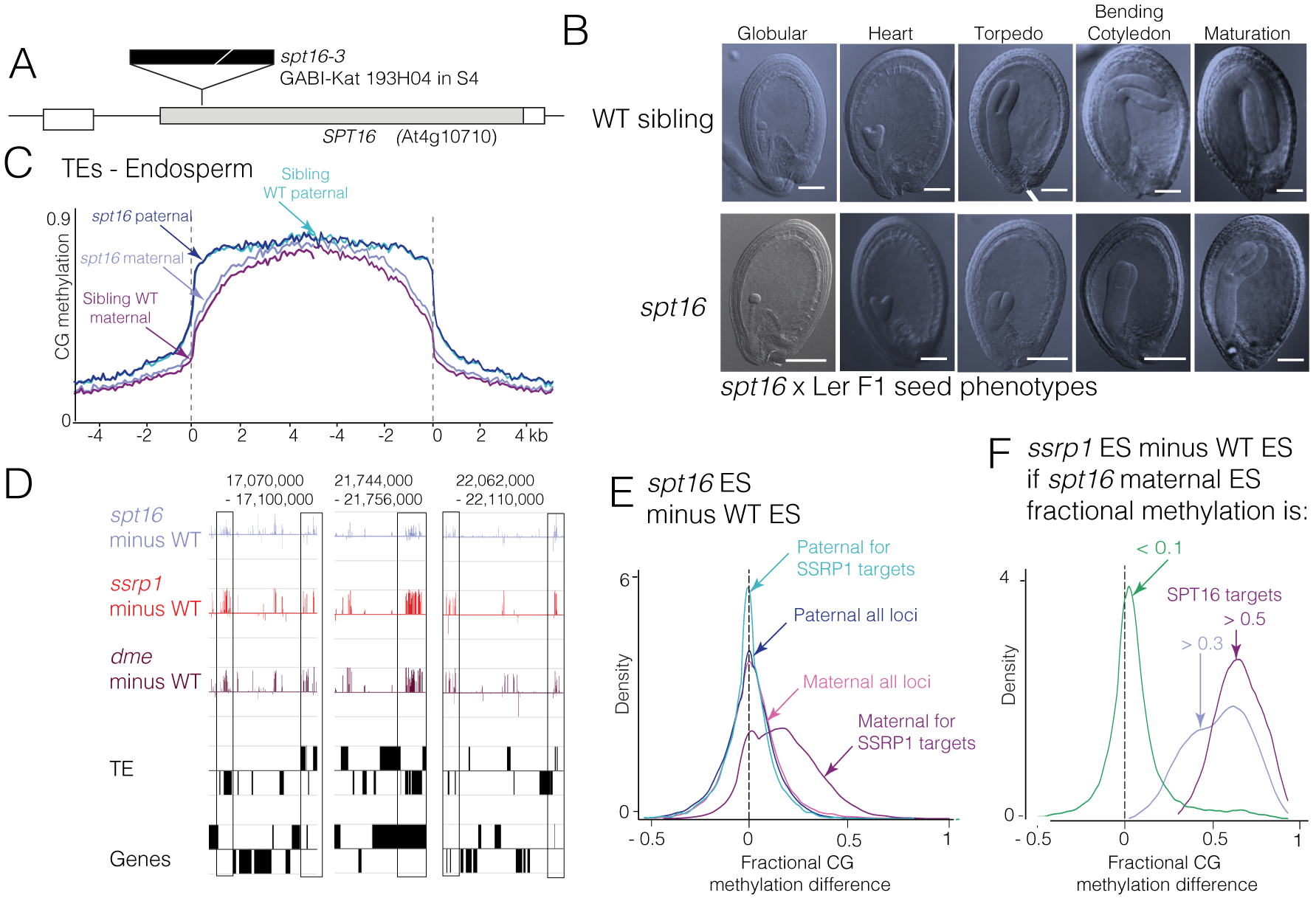
SPT16 contributes to DNA demethylation. (A) Diagram showing the *SPT16* gene structure and location of the GABI-kat 193H04 T-DNA insertion *spt16-3* (B) Photographs of developing F1 seeds from heterozygous *spt16-3* mutant plants crossed as females to wild-type L*er* pollen. ‘Delayed’ and ‘Normal’ seed fractions were used as mutant and wild-type control samples respectively. The morphological stages of wild-type embryos are indicated, scale bar = 100 um (C) Patterns of TE CG DNA methylation on *spt16-3* mutant ‘delayed’ maternal and paternal, and ‘normal’ WT-sibling (seed from same siliques) maternal and paternal endosperm alleles. Arabidopsis TEs were aligned at the 5’ or 3’ ends, and average methylation for all cytosines within each 50-bp interval is plotted. Dashed lines represent the points of alignment. (D) Genome browser alignments of DNA methylation with genes and TE annotations at selected loci of Arabidopsis Chromosome 1. Traces show CG methylation differences for mutant minus WT maternal endosperm in *spt16-3, ssrp1-3* and *dme-2* (9). Positive numbers indicate hypermethylation in the mutant genome. Regions of hypermethylation that overlapping in each mutant are boxed. (E) Kernel density plots of CG methylation differences between *spt16-3* mutant endosperm and WT endosperm for maternal and paternal alleles, for all sites and for SSRP1/FACT target loci, as defined in Figure 1C. (F) Kernel density plots of CG methylation differences between *ssrp1-3* mutant endosperm and WT endosperm, specifically for SPT16 targets, and for non-SPT16 target DNA. SPT16 CG targets are defined as those loci with a fractional CG methylation level of either > 0.3 or 0.5 in *spt16-3* mutant endosperm when compared to wild-type, and non-targets a level of < 0.1.

Plants heterozygous for *spt16-3* in the Columbia-0 (Col-0) ecotype were pollinated with wild-type pollen in the L*er* ecotype and hand-microdissected F1 embryo and endosperm were isolated 8-10 days after pollination (DAP). Next-generation bisulfite sequencing was carried out to compare DNA methylation in delayed (maternal *spt16-3* mutant) versus normally-developing (WT siblings) endosperm and embryos. Data were aligned according to the 5’ and 3’ ends of transposable elements (TEs) revealing very slight CG hypermethylation in *spt16-3* mutant maternal endosperm compared to wild-type (Figure 3C), whilst *spt16-3* embryo methylation was identical to wild type (Figure S3C). However, *spt16-3* maternal endosperm hypermethylation could be distinctly detected at SSRP1 and DME target loci (Figure 3D) and by kernel density analysis for only SSRP1 target loci, specifically on the maternal *spt16-3* endosperm allele (Figure 3E, Maternal for SSRP1 targets, positive purple trace). Moreover, by plotting the fractional methylation difference between *ssrp1-3* and WT maternal endosperm at only those loci where *spt16-3* maternal endosperm was hypermethylated (*spt16* minus WT fractional methylation >0.3 or >0.5 (Figure 3E) defined as SPT16 targets), we show that *spt16-3* hypermethylated loci are also sites of *ssrp1-3* hypermethylation (Figure 3F, blue and dark purple traces). Non-*spt16-3* hypermethylated loci (fractional methylation <0.1 (Figure 3E) defined as non-SPT16 targets) instead give a very different density curve (Figure 3F, green trace), centered on zero and thus enriched for sites that are not hypermethylated in *ssrp1-3* endosperm. Therefore, the effect of the *spt16-3* mutation on endosperm DNA methylation is qualitatively the same as *ssrp1-3*, but weaker. These results are consistent with a model of central cell demethylation by DME mediated by both subunits of FACT.

### FACT and DME proteins interact in the nucleus *in vivo*

To investigate the relationship between FACT and DME, we measured whether they interacted *in vivo*. We used the Bimolecular Fluorescence Complementation (BiFC) assay (32) to detect protein-protein interactions by expressing combinations of full length DME and the N- and C- termini of SSRP1 and SPT16 proteins, linked to portions of the Yellow Fluorescent Protein (YFP) in Arabidopsis leaf protoplasts. We observed frequent bright fluorescent signals of reconstituted YFP in the nucleus of cells expressing SSRP1-N and SPT16-C, as expected given that they form the FACT complex (Figure 4A; SSRP1-C and SPT16-N constructs displayed self-activity so were not included). We also observed bright, albeit fewer, fluorescent signals in the nucleus of cells expressing SPT16-C and DME (Figure 4B), indicating that these proteins are closely localized, possibly within the same macromolecular complex. YFP signals tended to overlap regions of intense chromatin staining, indicating that they were localized in heterochromatin (Figure S4A and B, Hoechst staining). We observed DME and SSRP1 fluorescent signals to be very infrequent and faint (Figure S4A), indicating that the more direct interaction occurs between DME and the larger SPT16 subunit of FACT. No reconstituted YFP signals were observed in protoplasts transfected with DME-YFP-C alone (Figure S4B) or with control protein LHP1 (Figure S4C). These results suggest that the FACT complex and DME are closely localized in the nucleus.

**Figure 4.**
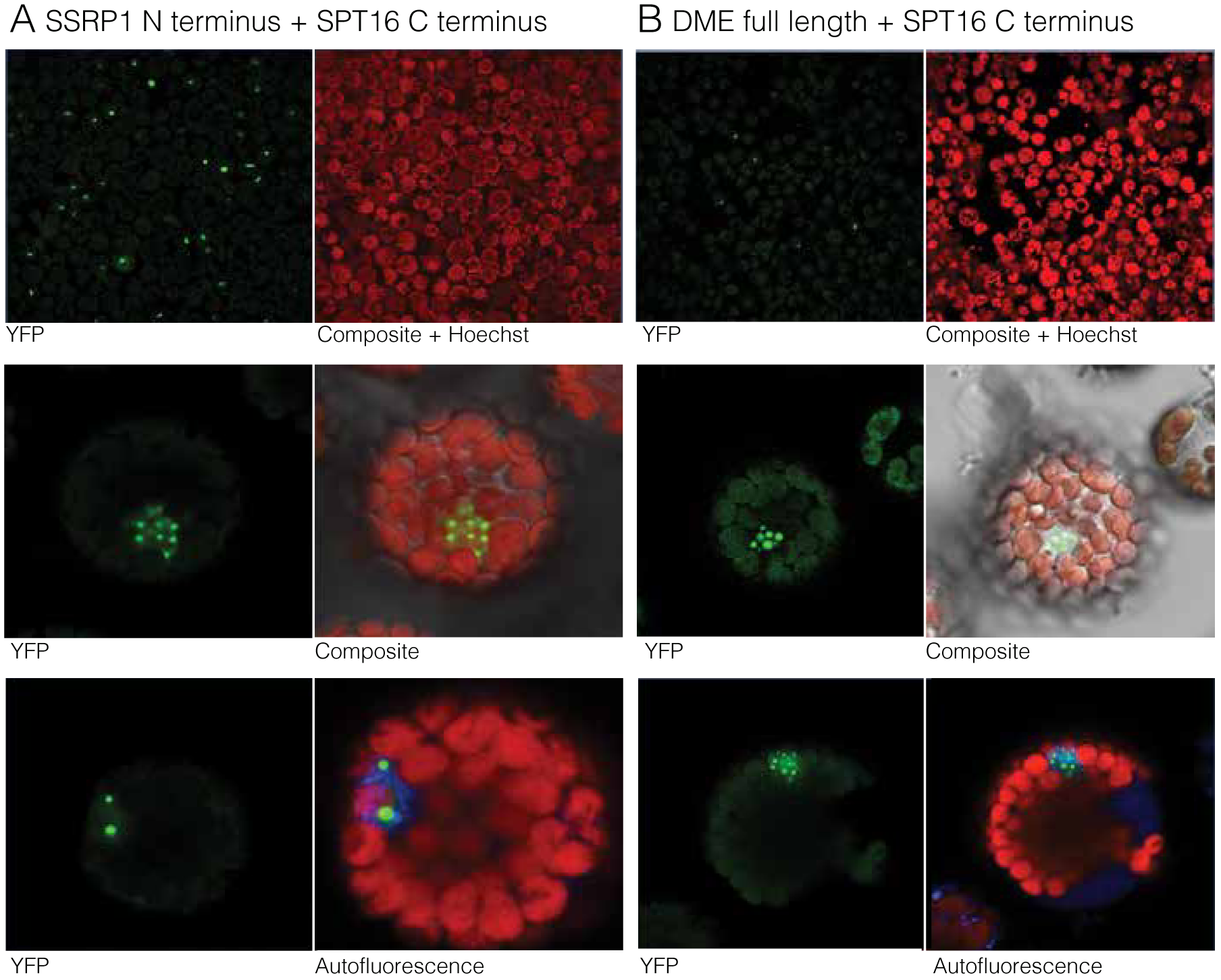
FACT and DME proteins interact in the nucleus in vivo. (A) Confocal fluorescence microscopy images from Bimolecular Florescence Complementation (BiFC) assays, showing the relative frequency of interactions as represented by YFP fluorescent signals generated by interactions between SSRP1-N terminal and SPT16-C terminal protein domains and (B) the SPT16-C terminal domain and DME protein. Top panels show lower magnification to demonstrate the frequency of interactions observed. Lower two panels show high magnification images show the location of fluorescent spots relative to the protoplast structure, and chromatin in the nucleus, stained with Hoechst 33342.

### FACT is required for regulation of a subset of imprinted genes

DME mediated demethylation in the central cell promotes both maternal and paternal expression of imprinted genes, since DNA methylation can either inhibit, or promote gene transcription depending on genomic context (33, 34). The short, euchromatic transposable elements overrepresented in the targets of DME, and representing a large proportion of FACT-DME targets, are often found upstream of imprinted genes (Ibarra et al. 2012). The SSRP1 subunit of FACT was previously observed to be required for demethylation of the SINE element controlling imprinted FWA expression, and for expression of the PRC2 subunit MEDEA (MEA) (26). To investigate the role of FACT in imprinted gene expression, we surveyed maternally and paternally expressed ‘stringent’ imprinted genes (35) and correlated them with either our genome-wide dataset of FACT target sites, or with all Arabidopsis genes, using ends-analysis (Figure 5A). Similar to DME-only target sites (Ibarra et al., 2012) FACT-DME shared target sites were significantly enriched compared to all genes (by Fisher exact test) at the 5’ end and around the transcriptional start and termination sites of maternally expressed imprinted gene loci, indicating that the FACT-dependent subset of DME-target sites do include imprinted regions (Figure 5A). FACT-DME targets were also significantly enriched in the gene body of maternally expressed genes, but not at the 3’ end. FACT-DME targets were not significantly associated with paternally expressed imprinted genes. To look at individual imprinted gene loci, we analyzed DNA methylation in wild-type, *ssrp1-3, spt16-3* and *dme-2* mutant maternal endosperm for both maternally and paternally expressed imprinted genes locus-specifically (Figure 5B and C). Methylation of imprinting control regions for maternally expressed imprinted genes is aberrant in *ssrp1-3 and spt16-3* endosperm (Figure 5B), consistent with the contribution of FACT to imprinted gene regulation, as a function of facilitating access of DME to demethylate heterochromatic DNA. Some key imprinted genes (Figure 5C), such as FIS2 which result in seed abortion when not expressed (36), do not appear to be regulated by FACT, consistent with the reduced seed abortion in *ssrp1-3* and lack of seed abortion in *spt16-3* mutant siliques compared to *dme-2* (Figure 5C).

**Figure 5.**
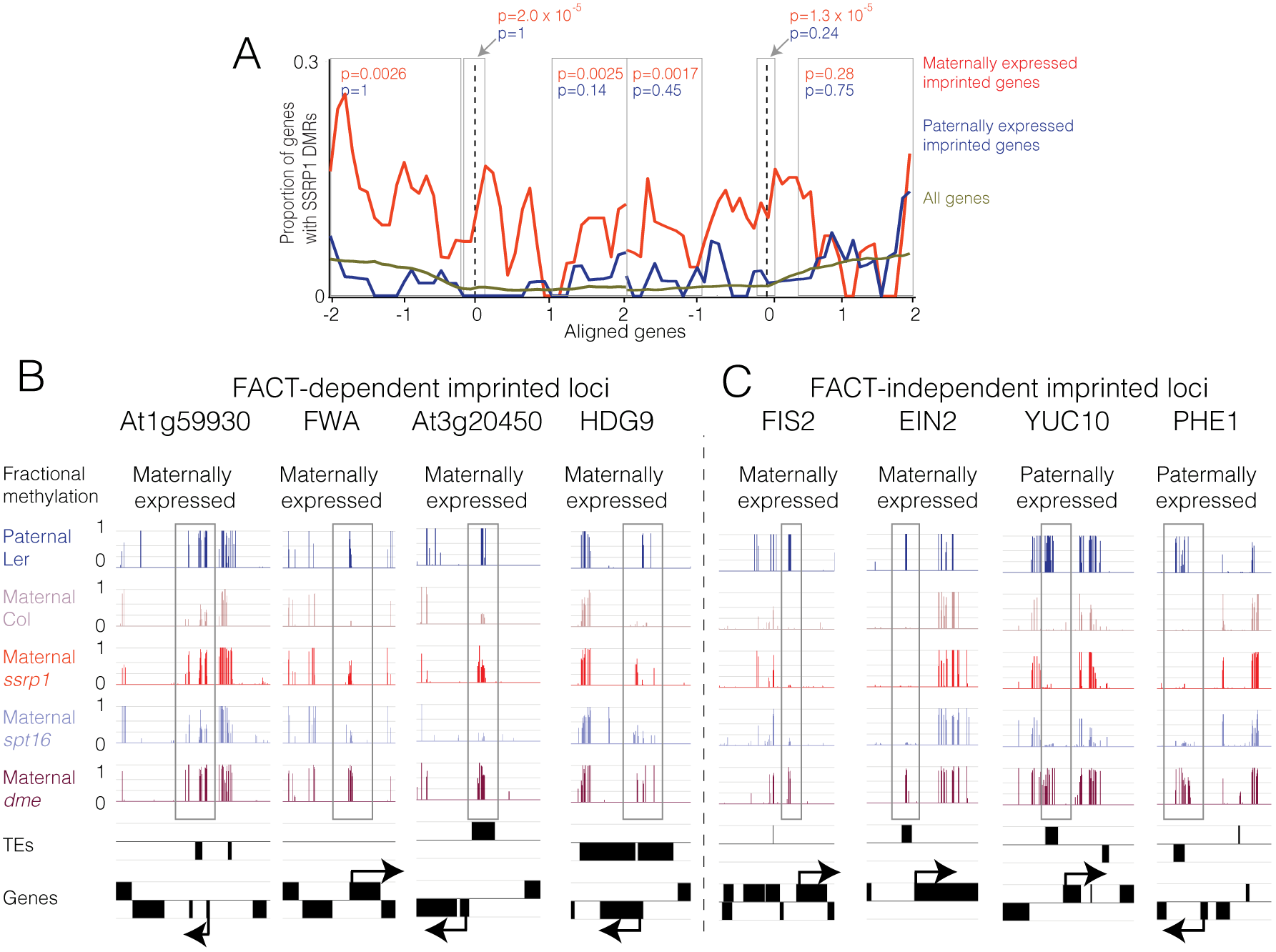
FACT is required for regulation of a subset of imprinted genes. (A) The distribution of significantly differentially methylated regions (DMRs, as in Figure 2B) between *ssrp1-3* and WT maternal endosperm near to genes. Genes were aligned at the 5’ end (left dashed line) or the 3’ end (right dashed line) and the proportion of genes with DMRs in each 100-bp interval is plotted. DMR distribution is shown with respect to maternally expressed imprinted genes (red trace), paternally expressed imprinted genes (blue) and all genes (brown trace). Significance of DMR enrichment with respect to all genes (Fisher’s exact test) for particular genic regions is shown in gray boxes. Arabidopsis imprinted genes were collated from (35). (B) Snapshots of CG methylation in endosperm near imprinted genes that were dependent on FACT regulation (gained hypermethylation in *ssrp1-3* mutant endosperm) and (C) independent of FACT regulation (unaffected by *ssrp1-3* mutation). Paternal (L*er*) endosperm is in blue, with maternal alleles of the *ssrp1-3* mutant in red*, spt16* mutant in light blue, *dme* mutant in maroon, and maternal wild-type Col-0 in pink, aligned to annotated genes and TEs.

### FACT is required for DME-mediated demethylation in long TEs enriched with H3K9me2

To assess the characteristics of DME targets that require FACT for DNA demethylation, we aligned genome-wide *ssrp1-3* endosperm methylome data alongside *dme-2* endosperm methylome data (9) according to the 5’ and 3’ ends of TEs (Figure 6A) and genes (Figure S6A). In the bodies of aligned TEs (away from the points of alignment), hypermethylation in *ssrp1-3* mutant maternal endosperm is as high as that of the paternal endosperm allele, which is not demethylated, and of the maternal *dme-2* mutant. This indicates that in large TE bodies, FACT is always required for DME access. In agreement with this, whilst for both DME-only and DME-FACT shared sites, smaller TEs are the most prevalent target, TE length is positively correlated with FACT-DME shared targets compared to DME-only targets (Figure S6B). In TEs above 1 and 2 kb, there remains a difference in methylation at the TE edge between *ssrp1-3* and *dme*, (Figure S6C and D) but in TEs above 4 kb, the difference in methylation between *dme-2* and *ssrp1-3* is lost (Figure 6B and C). Long TEs are enriched in pericentromeric DNA and when we calculated the relative enrichment of FACT-DME shared and DME-only differentially methylated regions across the genome, FACT-DME shared sights were highly enriched at pericentromeric regions, whereas DME-only sites were more frequent than FACT-DME shared sites in chromosome arms (Figure 6D).

**Figure 6.**
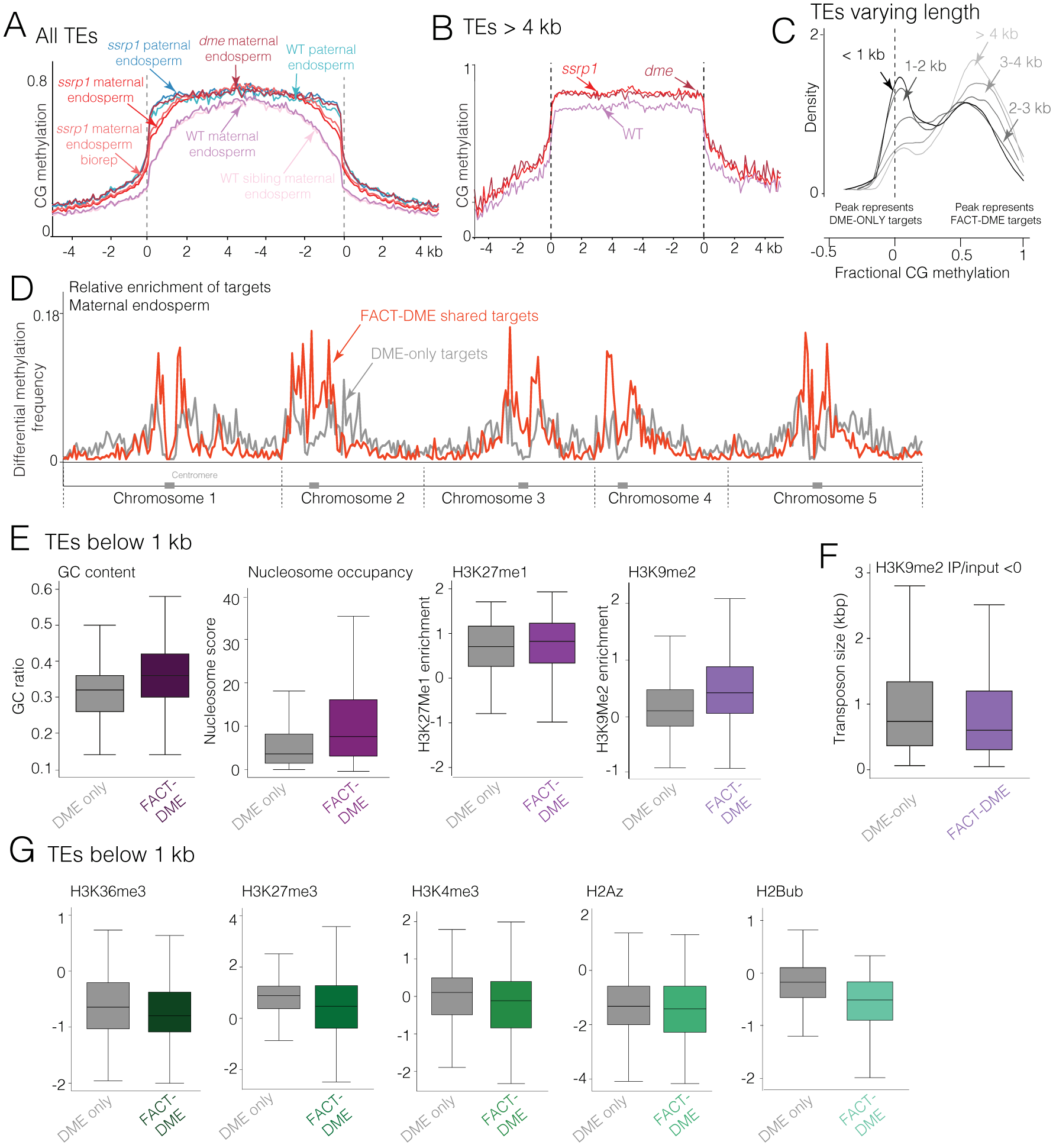
FACT is required for DME-mediated demethylation in heterochromatin but not euchromatin. (A) Average CG methylation in TEs in wild-type maternal and paternal, *ssrp1-3* mutant maternal (two biological replicates) and paternal, wild-type sibling maternal (from same siliques as *ssrp1-3* mutant) and *dme-2* maternal endosperm genomes is plotted with aligned 5’ and 3’ ends, as in Figure 2. (B) Average CG methylation in TEs longer than 4 kb, in wild-type, *ssrp1-3* and *dme-2* maternal endosperm genomes. (C) Kernel density plot of the methylation status of *dme-2* hypermethylated loci (fractional methylation difference compared to wild-type of >0.5) in *ssrp1-3* mutant endosperm, as in Figure 2A, for loci grouped by TE length (<1 kb, 1-2 kb, 2-3 kb, >4 kb). (D) Relative enrichment of FACT-DME shared and DME-only differentially methylated regions (defined as >0.5 fractional methylation difference for both, or >0.5 for DME and <0.1 for SSRP1, respectively, including at least 20 sequenced cytosines) in 300 kb intervals across the Arabidopsis genome. Increased density of methylation differences correlates with pericentromeric regions. (E) Box plots showing the relative enrichment of GC content, nucleosome occupancy and (log_2_ IP/INPUT) of H3K27me1 and H3K9me2 modifications between DME-only CG endosperm targets and FACT-DME shared endosperm targets within 50 bp windows, in TEs <1kb. (F) Box plot showing distribution of TE sizes across DME-only CG endosperm targets and FACT-DME shared endosperm targets within 50bp windows for regions with low H3K9me2 enrichment (log_2_ IP/INPUT <0) (G) Box plots showing relative enrichment (log_2_ IP/INPUT) of H3K36me3, H3K27me3, H3K4me3, H2AZ and H2Bub in DME-only and FACT-DME shared targets as for (E).

In short TEs and TE edges, *ssrp1-3* hypermethylation is less severe than in the *dme-2* mutant (Figure 6A), thus FACT is only required for demethylation at a subset of DME short TE targets. DME homologs ROS1, DML2, and DML3, which act in sporophytic tissues to ‘prune’ DNA methylation at certain sites, require histone acetyltransferase INCREASED DNA METHYLATION 1 (IDM1) and the IDM2 α-crystallin domain protein to gain access to DNA at a subset (10 %) of sites in regions depleted in H3K4me2 (37, 38). To determine whether chromatin structure similarly dictated the requirement for FACT in certain short TEs, we correlated the groups of DME-only and FACT-DME targets with structural chromatin features in aerial plant tissues (39–43), in TEs below 1 kb. We observed that FACT is more frequently required for DME activity in GC-rich regions with high nucleosome occupancy and increased levels of compact chromatin markers H3K27me1 and H3K9me2 (Figure 6E) (44). This is consistent with the requirement for FACT in longer TEs, which tend to have a heterochromatic structure (45, 46). In fact, by analyzing the enrichment of DME-only and FACT-DME shared targets according to TE length, in only those TEs with low H3K9me2 occupancy, the positive relationship between FACT-DME targets and increased TE length is lost (Figure 6F). DME-only target TEs below 1 kb tended to be enriched with markers of open chromatin; H3K27me3, H3K36me3, H3K4me3, H2AZ and, most strikingly, H2Bub (Figure 6G). Thus, FACT is required for DME access to all long TEs since they tend to be located in relatively heterochromatic DNA, and are enriched with histone modifications such as H3K9me2. Conversely, FACT is variably required for DME access to short TEs, which are more likely to occur in open chromatin, depending on the specific chromatin structure of those TEs, whereby at relatively euchromatic TEs (i.e. those with less H3K9me2), DME does not require FACT.

### H1 presence mediates the requirement for FACT at certain DME-target loci

The SSRP1 subunit of FACT is an HMG domain protein, which are known to compete with the histone linker H1 for chromatin occupancy (47). H1 binds to the nucleosome core and is strongly associated with heterochromatin (48). In addition, its presence is known to impede DNA accessibility in both euchromatin and heterochromatin (49). H3K9me2 enrichment, which we show to often be present at loci where FACT-is required for DME activity, is also correlated with regions of H1 occupancy, so we sought to determine whether H1 may impede access of DME in a manner that contributes to the requirement for FACT for DME activity

Using plants homozygous for *h1.1* and *h1.2* alleles (46), referred to as homozygous *h1*, we generated mutant plants that were also heterozygous for *ssrp1-3*. We did not observe any rescue of the *ssrp1-3* seed delay and abortion phenotype (Table S3), indicating that H1 does not wholly dictate the need for FACT in DME activity. To determine if H1 plays a more modest role, we pollinated plants homozygous for *h1* and heterozygous for *ssrp1-3* in the Col-0 ecotype with wild-type pollen in the L*er* ecotype. We then analyzed the maternal methylomes of F1 microdissected endosperm from delayed F1 seeds (maternal mutant *h1 ssrp1-3) versus* their normally developing siblings (maternal mutant *h1*), and compared them to maternal mutant *ssrp1-3* endosperm *versus* wild-type endosperm. We could not detect an obvious decrease in maternal genome-wide hypermethylation in *h1 ssrp1-3* compared to *ssrp1-3* (Figure 7A). However, by looking specifically at all DME-target loci both genome-wide and at individual FACT-DME targets (Figure 7B and C, respectively), we identified a decrease in hypermethylation in *h1 ssrp1-3* compared to *ssrp1-3*, specifically at FACT-DME shared targets, consistent with an effect of H1 on FACT-DME targets, but not DME-targets (Figure 7B, C and D). *h1* mutant seedling DNA is hypomethylated at euchromatic TEs genome-wide (46); thus, it is possible that the loss of hypermethylation seen in our triple mutant was simply due to an underlying absence of DNA methylation in *h1* mutant endosperm. We analyzed *h1* mutant seedling DNA methylation specifically at loci that are differentially methylated between *h1 ssrp1* and *ssrp1* mutant maternal endosperm ‘FACT-DME DMRs associated with H1’ (n=565, Figure 7D, Table S1). *h1* mutant seedlings were not predominately hypomethylated at FACT-DME DMRs associated with H1 (Figure 7C, and E), indicating that the lack of hypermethylation in *h1 ssrp1* mutants at FACT-DME targets is due the *h1 ssrp1* genotype rather than a hypomethylated *h1* background. Thus, loss of H1 partially suppressed the hypermethylation caused by the *ssrp1-3* mutant, indicating that H1 may impede DME access to chromatin, which is relieved by FACT.

**Figure 7.**
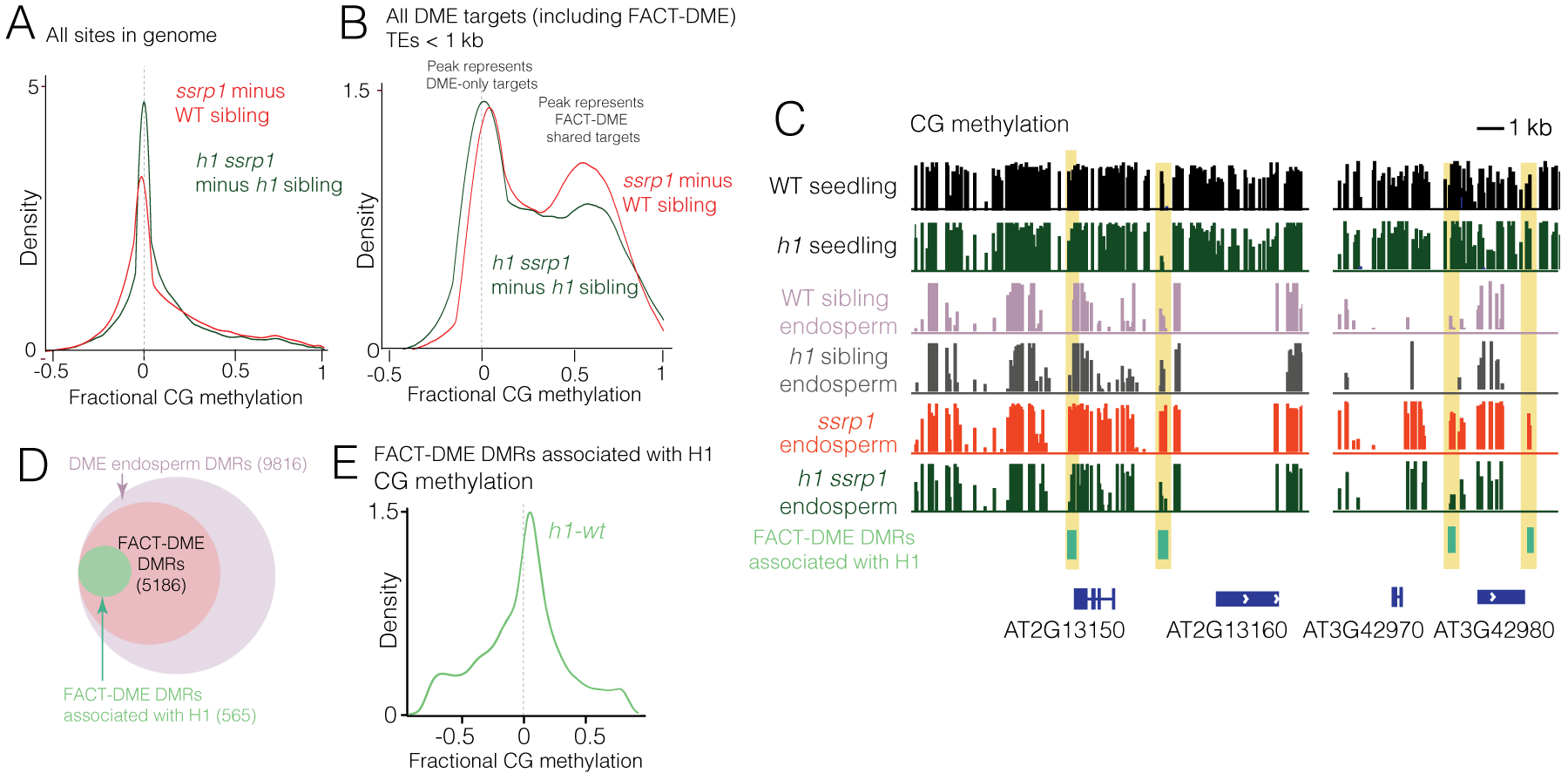
H1 presence mediates the requirement for FACT at certain DME-target loci. (A) Kernel density plot of *h1.1/h1.2/ssrp1-3* (*h1 ssrp1*) triple mutant endosperm (delayed) minus *h1.1/h1.2/*SSRP1 (*h1* sibling; normal development) endosperm maternal methylation, compared to *ssrp1-3* (delayed) minus SSRP1 (WT sibling, normal development) maternal endosperm methylation, genome-wide and (B) for DME targets only, in short TEs (<1 kb). (C) CG methylation profiles for WT and *h1* seedlings, and WT, *h1* sibling, *ssrp1* and *h1 ssrp1* mutant endosperm at examples of FACT-DME DMRs associated with H1. (D) Venn diagram to illustrate proportion of FACT-DME DMRs that are associated with H1 occupancy (565 DMRs, >20 % methylation difference between *h1 ssrp1* and *ssrp1* mutant endosperm, p<0.001). (E) Kernel density plot of *h1* seedling (46) minus WT seedling CG methylation, plotting only FACT-DME DMRs associated with H1 where *ssrp1 h1* was hypomethylated compared to *ssrp1,* showing that DNA methylation in these regions is largely the same in WT and *h1* knockout plants.

## Discussion

Like ATP-dependent chromatin remodeling enzymes, FACT dramatically increases the accessibility of nucleosomal DNA. Current evidence suggests that FACT interaction with the nucleosome promotes formation, or stabilization, of a looser structure, still bound to DNA, but more likely to undergo reorganization through H2A/H2B displacement (20, 50, 51). Our data show that FACT is required for DME-mediated demethylation in regions with high nucleosome occupancy, and enrichment for posttranslational histone modifications associated with heterochromatin, particularly H3K9me2 (Figure 6D, E, and F). Less than half of DME target sites are accessible without FACT involvement (Figure 2A and B). These ‘DME-only’ sites were shorter TEs, enriched with euchromatic markers such as H2Bub (Figure 6G and S6D), and depleted in H3K9me2, thus representing loci with chromatin that was more accessible to proteins such as DME.

Genomic loci exhibiting enrichment of H3K36me3 and H2Bub, at least in seedling tissues, were anti-correlated with sites of FACT activity in DME-mediated DNA demethylation as observed in endosperm (Figure 6G). This is striking because during transcription, H2Bub is a positive regulator of FACT (52). Similarly, methylation of H3K36 mediates the requirement for FACT in transcription initiation, and H2BK123/120Ub1 stimulates FACT activity in transcriptional elongation and nucleosome reassembly following transcription (52–54). These data indicate that the mode of FACT action with DME DNA glycosylase during reproduction differs from that during transcription. This is reminiscent of recent data from experiments in human cells, showing that upon oxidative stress, FACT is relocated away from transcribed DNA to regions requiring repair, where it remodels chromatin to promote base excision repair, facilitating the interaction between 8-Oxoguanine glycosylase (OGG1) and DNA (55).

A further key feature of heterochromatin is the association of nucleosomes with linker histone H1. FACT directly interacts with H1 *in vitro* and *in vivo* in mammalian cells (56, 57), and the SSRP1 FACT subunit has an HMG domain, which tend to compete with H1 molecules for nucleosome binding, weakening the interaction between H1 and chromatin (58). Thus, FACT may chaperone H1-containing nucleosomes in Arabidopsis, contributing to the enhancement of DNA accessibility for DME activity during reproduction. Removing H1 did not obviate the need for FACT in DME activity (Figure 7A); however, we found a decrease in hypermethylation specifically at FACT-DME target sites in mutant homozygous *h1,* maternal *ssrp1* endosperm (Figure 7B, C and D), corresponding to approximately 10 % of FACT-DME target sites. This is reminiscent of chromatin-dependent mechanisms of DNA methylation (4, 46). Notably, chromatin remodeler DEFICIENT IN DNA METHYLATION 1 (DDM1), facilitates access of DNA methyltransferases to H1-bound chromatin (46). These data are consistent with a gradient of heterochromatic status (46) within the Arabidopsis central cell genome, whereby DME targets range from euchromatic, and accessible, to more heterochromatic, where chromatin accessibility decreases below a threshold, at which point FACT is required.

The requirement for FACT in DME activity in the central cell, but not the vegetative cell, is intriguing (Figure 2C). Since the vegetative cell nucleus is separated from its somatic precursor by only one cell division, it is possible that SSRP1 or SPT16 proteins are still present in this tissue, contributing to DNA demethylation and thus masking any molecular phenotype in *ssrp1-3* mutant pollen. Alternatively, the explanation may involve chromatin. Although vegetative and central cell nuclei are relatively decondensed (59), they have highly diverse fates; the vegetative cell undergoes no further division and is a terminally differentiated cell, whereas the central cell is fertilized and goes on to form the endosperm. It is likely the vegetative and central cells’ chromatin conformation will be different, which may explain why FACT is apparently not required for DME activity in pollen (60), but is needed in the central cell.

## Experimental Procedures

### Arabidopsis mutants

*ssrp1-3* (26) and *h1.1/h1.2* double mutants (46) were as described previously. The *spt16-3* T-DNA insertion line (GK_193H04) was obtained from the GABI-Kat collection (31) at the Nottingham Arabidopsis Stock Centre (NASC; (61)). All mutants were in the Columbia-0 (Col-0) background. T-DNA insertions were confirmed by PCR and the location of *spt16-3* identified using Sanger sequencing by the Barker Hall Sequencing facility at UC Berkeley.

### Isolation of Arabidopsis endosperm and embryos

Wild-type Col-0 and mutant Arabidopsis flower buds were emasculated at flower stage 12-13 using fine forceps and pollinated with L*er* pollen 48 hours later. Eight to ten days after pollination (DAP) developing F1 seeds (torpedo to bending cotyledon stage) were immersed in dissection solution (filter-sterilized 0.3 M sorbitol and 5 mM pH 5.7 MES) on sticky tape and dissected by hand under a stereo-microscope using fine forceps (Fine Science Tools, Inox Dumont #5) and insect mounting pins. The seed coat was discarded, and debris removed by washing collected embryos or endosperm five to six times with dissection solution under the microscope. For *ssrp1-3* and *spt16-3* mutant normal and delayed sibling seed collection, seeds were divided into fractions according to embryo development. At 8 DAP, delayed seeds tended to be at the torpedo to early-linear cotyledon stages, whereas normally developing seeds were late-linear cotyledon to bending cotyledon stages (Figure 1B and 3B). Seeds with an intermediate developmental stage were discarded.

### Delayed- and normally-developing sibling seed genotyping

Aliquots of dissected endosperm gDNA from delayed and normally developing sibling seeds (i.e. isolated from within the same siliques), used subsequently to generate bisulfite sequencing libraries, were used as templates for genotyping. *ssrp1-3* and *spt16-3* genotyping amplicons were generated using primers flanking the *ssrp1-3* SNP or the T-DNA insertion in *spt16-3*. *ssrp1-3* amplicons were cloned into TOPO TA, colony PCR and Sanger sequencing performed on at least 50 individual strands, and ratios of WT and mutant alleles calculated. *spt16-3* genotyping was carried out using gel electrophoresis to separate differentially sized PCR products.

### Isolation of vegetative cell and sperm nuclei

Pollen was isolated from wild-type (Col-0) and *ssrp1-3* heterozygous plants as described previously (62, 63). Vegetative cell and sperm nuclei were extracted from mature pollen and fractionated by fluorescence activated cell sorting as described previously (62, 63).

### Bisulfite sequencing library construction

As described previously, genomic DNA was isolated from vegetative cell and sperm nuclei (63), endosperm, and embryo (18). Single-end bisulfite sequencing libraries for Illumina sequencing were constructed as in (18) with minor modifications. In brief, about 50 ng of genomic DNA was fragmented by sonication, end repaired and ligated to custom-synthesized methylated adapters (Eurofins MWG Operon) according to the manufacturer’s instructions for gDNA library construction (Illumina). Adaptor-ligated libraries were subjected to two successive treatments of sodium bisulfite conversion using the EpiTect Bisulfite kit (Qiagen) as outlined in the manufacturer’s instructions. The bisulfite-converted library was split between two 50 ul reactions and PCR amplified using the following conditions: 2.5 U of ExTaq DNA polymerase (Takara Bio), 5 μl of 10X Extaq reaction buffer, 25 μM dNTPs, 1 μl Primer 1.1 and 1 μl multiplexed indexing primer. PCR reactions were carried out as follows: 95°C for 3 minutes, then 14-16 cycles of 95 °C 30 s, 65 °C 30 s and 72 °C 60 s. Enriched libraries were purified twice with AMPure beads (Beckman Coulter) prior to quantification with the Qubit fluorometer (Thermo Scientific) and quality assessment using the DNA Bioanalyzer high sensitivity DNA assay (Agilent). Sequencing on either the Illumina HiSeq 2000/2500 or HiSeq 4000 platforms was performed at the Vincent J. Coates Genomic Sequencing Laboratory at UC Berkeley.

### Bisulfite data analysis

Sequenced reads were sorted and mapped to the TAIR8 or TAIR10 Col-0 and L*er* genomes as described previously (34). Gene and TE ends analysis, box plots and kernel density plots were generated as previously described (9), using only windows with at least 20 informative sequenced cytosines, and fractional methylation of at least 0.7 (CG), 0.4 (CHG) or 0.08 (CHH) in at least one of the samples being compared. Differentially methylated regions (DMRs) in *ssrp1-3* endosperm and vegetative cell were also generated as previously (9), whereby windows with a fractional CG methylation difference of at least 0.3 between *ssrp1-3* endosperm/vegetative cell and WT (Fisher’s exact test p-value < 0.001 for endosperm, <10^-10^ for vegetative cell) and were merged to generate larger DMRs if they occurred within 300 bp. DMRs were retained for further analysis if the fractional CG methylation across the whole DMR was 0.3 greater in *ssrp1* endosperm than in wild-type endosperm (Fisher’s exact test p-value < 10^-10^), and if the DMR was at least 100 bp. FACT DMRs associated with H1 occupancy were a subset of FACT (ES) DMRs, and had a fractional methylation difference of at least 0.2 between *h1.1 h1.2 ssrp1-3* endosperm and *ssrp1-3* endosperm, Fisher’s exact test p-value <0.001. DMRs overlapping with imprinted gene loci were identified as for (9), and using unsorted reads, except that the list of imprinted genes was obtained from (35).

### Correlations of histone modifications and genomic attributes

Nucleosome enrichment data was from (39), H3K9me2 from (40), H3K27me3 from (41), H2A variants from (42) and other modifications from (43).

### Bimolecular Fluorescence and confocal microscopy

To analyze the interaction between the DME protein and the FACT complex in vivo, full-length *cDME* (5.2 kb of the *At5g04560.*2 transcript) was cloned into a *pSAT4-nEYFP-C1* vector. Both C-terminal *cSPT16* (1.9 kb) and N-terminal *cSSRP1* (1.9 kb) were cloned into a *pSAT4-cEYFP-C1-B* vector. Pairs of constructs were introduced into Arabidopsis leaf protoplasts by PEG transfection as described previously (64). After incubation, fluorescence was observed using the Zeiss Confocal Laser Scanning Microscope LSM700.

## Author Contributions

Conceptualization, J.M.F., T.K., Y.C., D.Z., and R.L.F.; Methodology, J.M.F., M.Y.K., G.T.P., P.H.H., J.C., Y.I.; Formal Analysis, J.M.F., M.Y.K., P.H.H., J.C. Investigation, J.M.F., M.Y.K., G.T.P., P.H.H., M.N., J.C., S.L. and Y.I.; Visualization, J.M.F., G.T.P., and H.Y.; Writing – Original Draft, J.M.F.; Writing – Review & Editing, J.M.F., Y.C., Y.I., T.K., M.Y.K., S.L., M.N., D.Z. and R.L.F.; Funding Acquisition, T.K., Y.C., D.Z. and R.L.F.; Resources, T.K., Y.C., D.Z. and R.L.F. Supervision, Y.C., T.K., D.Z. and R.L.F.

## Acknowledgements

The authors would like to thank Christina Wistrom for her management of the UC Berkeley Oxford Tract greenhouse facility. This work used the Vincent J. Coates Genomics Sequencing Laboratory at UC Berkeley, supported by NIH S10 Instrumentation Grants S10RR029668, S10RR027303 and S10OD018174, and the authors would like to particularly thank Shana McDevitt for her assistance. This work was funded by NSF (IOS-1025890) and NIH (R01-GM069415) grants to D.Z and R.L.F., by NRF of Korea (2017R1A2B2007067) and the Next-Generation BioGreen 21 Program (PJ011018) grants to Y.C., and by KAKENHI (JP16H06471) to T.K. The authors declare no conflicts of interest.

